# Data Independent Acquisition Mass Spectrometry of the Human Lens Enhances Spatiotemporal Measurement of Fiber Cell Aging

**DOI:** 10.1101/2021.05.13.444062

**Authors:** Lee S Cantrell, Kevin L Schey

## Abstract

The ocular lens proteome undergoes post-translational and progressive degradation as fiber cells age. The oldest fiber cells and the proteins therein are present at birth and are retained through death. Transparency of the lens is maintained in part by the high abundance crystallin family proteins (up to 300 mg/mL), which establishes a high dynamic range of protein abundance. As a result, previous Data Dependent Analysis (DDA) measurements of the lens proteome are less equipped to identify the lowest abundance proteins. In an attempt to probe more deeply into the lens proteome, we measured the insoluble lens proteome of an 18-year-old human with DDA and newer Data Independent Analysis (DIA) methods. By applying library free DIA search methods, 4,564 protein groups, 48,474 peptides and 5,577 deamidation sites were detected: significantly outperforming the quantity of identifications in using DDA and Pan-Human DIA library searches. Finally, by segmenting the lens into multiple fiber cell-age related regions, we uncovered cell-age resolved changes in proteome composition and putative function.

## Introduction

The ocular lens is a transparent tissue responsible for transmission and focusing of light to the retina. Unlike the majority of all tissue, lens fiber cells are not degraded after formation and organelles degrade in early cellular maturation, leading to cessation of protein synthesis. Since genetic material is degraded, proteomic measurement is exclusively capable of characterizing each stage of cellular maturation. Age-related measurements of lens fiber cell proteomes are facilitated by the inherent spatiotemporal gradient of the lens wherein fibers of the innermost nucleus of the lens are as old as the subject and surrounding concentric layers of fibers are progressively younger. Here, we divide the lens into three age-related regions: the inner nucleus that is composed of fiber cells formed *in utero* and infancy, the outer nucleus that contains fibers differentiated in early childhood, and the cortex that corresponds to fibers differentiated in early adulthood (Figure 1).

**Figure 1.**
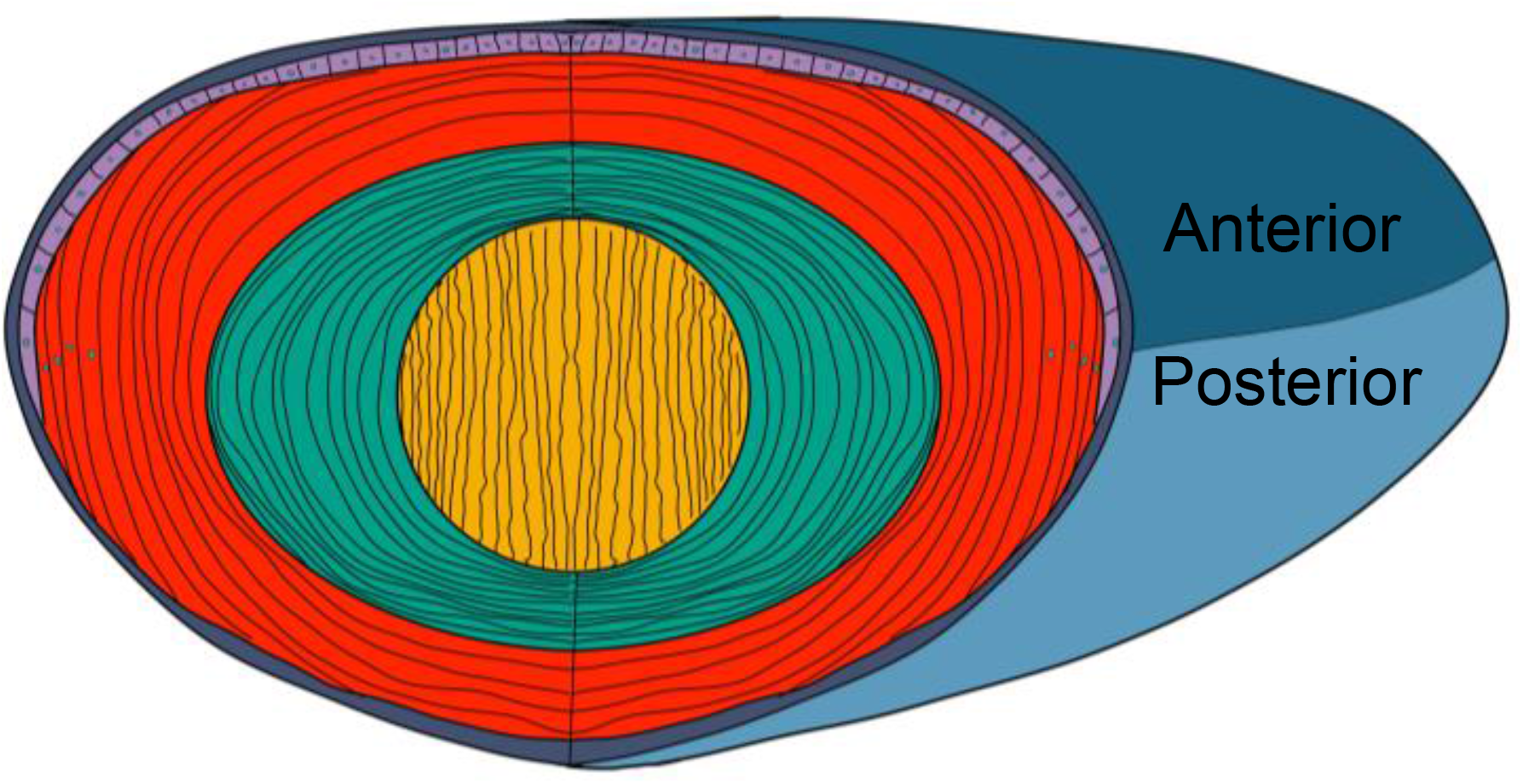
Cartoon of lens with regions annotated. Fiber cells are differentiated from epithelial cells (purple) into the cortex (red) where organelles (teal circles) degrade. After the fifth decade of life, an extracellular diffusion barrier forms between the outer nucleus (green) and cortex, and the oldest fibers, formed in utero may be found in the inner nucleus (orange). The entire lens is encompassed by a collagenous capsule (blue).

Throughout life, transparency of the lens must be maintained to prevent cataract.^1^ In the absence of vasculature, metabolites are hypothesized to convect through the lens by a microcirculation system (MCS) where metabolite influx at the poles proceeds towards the inner nucleus and outflow proceeds through the equatorial region of the lens. In the proposed system, flux is established by a sodium gradient near the equatorial epithelium and water is transported through membranes by aquaporin and gap junction proteins. The MCS is critical for the delivery of glutathione to the inner nucleus of the lens, where oxidative stress may otherwise induce post-translational modifications (PTMs) that modify protein structure, function and induce light scattering by protein aggregation. In addition to glutathione conjugation to oxidized species, α-crystallin contributes to aggregation prevention by binding to partially unfolded substrates as a small heat shock protein.^2,3^ After the fifth decade of life, a barrier to extracellular metabolite transport is established, further challenging maintenance of lens transparency.^4,5^

Historically, measurement of the lens proteome and its post-translational modifications have centered on the crystallin protein family due to their overwhelming abundance in the lens: estimated as high as 90% by mass and 3,000 mg/mL.^6–9^ The high concentration of lens crystallins reduces the technical ability to measure low abundance proteins in the lens, especially with intensity biased methods such as Data Dependent Acquisition (DDA) MS,^10^ making the lens and excellent test tissue for DIA methodology. Further, biological insights may be gained by solubility-fractionation to deplete soluble crystallin proteins, isolating insolubilized proteins, cytoskeletal components, or proteins localized to the plasma membrane – like those in the MCS.^11,12^ Using a 13-pulse MuDPIT separation coupled to a LTQ Velos linear ion trap mass spectrometer, solubility fractionation provided previous identification of 951 human lens protein groups.^11^ More recently, 5,466 protein groups from young mouse lenses have been identified with a 9-step basic reverse phase (BRP) pre-separation and acquisition on an newer orbitrap instrument. For either mass analyzer, MuDPIT/BRP improves the quantity of protein and peptide identifications by approximately 400% over single-dimension analyses.^13,14^ While 2D separation is effective in increasing protein identifications, it is less quantitatively reproducible and acquisition times are significantly greater per samples.^15^ Further, if the lens is to be segmented into multiple fiber cell-age related regions for measurement, 2D separations become significantly more instrument intensive than single-dimension analyses.

Arguably the greatest limitation of DDA is its weakness for measuring low abundance peptides due to intensity biased selection of precursors.^16,17^ Alternatively, Data Independent Acquisition (DIA) methods including Sequential Window Acquisition of All Theoretical Mass Spectra (SWATH-MS), circumvent biased precursor selection by co-isolating all peptides within a small mass window along the LC gradient.^10^ The major limitation of DIA is data-processing where a spectral library is required to deconvolute individual product spectra which may represent multiple co-eluting precursors.^18^ Conventional spectral libraries are developed by combining a fraction of each sample in the cohort and sampling with DDA. To improve spectral library size, 2D separations and gas-phase fractionation are frequently employed. Constructing a library is a laborious and instrument intensive process, so widely applicable libraries such as the pan-human SWATH-MS library may be used instead.^19^ Alternatively, recent software developments include *in silico* library preparation.^20,21^ Here, we examine the application of DIA to multiple age-related regions of the human lens and discuss the impact that DIA, pre-developed libraries, and *in silico* searches have on annotating age-related changes in lens fiber cells.

## Materials and Methods

### Materials

An 18-year-old frozen human lens was obtained from NDRI (Philadelphia, PA). Sequence-grade modified trypsin was obtained from Promega (Madison, WI). S-Trap columns were purchased from Protifi (Farmingdale, NY). All other chemicals were purchased from Fisher Scientific (Waltham, MA).

### Urea Insoluble Protein Preparation

An 18-year-old, cataract free human lens (9.2 mm x 7.0 mm) was mounted with its equatorial axis parallel to the cryostat chuck before removal of the anterior and posterior poles of the lens yielding a 4.5 mm thick equatorial lens section. Concentric biopsy centerpunches were taken at 4.5- and 7-mm diameter to yield inner nucleus (0-4.5 mm), outer nucleus (4.5 – 7.0 mm) and cortex (7.0 – 9.2 mm) samples. Tissue was homogenized in buffer containing 25 mM Tris (pH8), 5 mM EDTA, 1 mM DTT, 150 mM NaCl, 1 mM PMSF. After homogenization, samples were centrifuged at 100,000g for 30 minutes and the supernatant discarded. Pellets were washed twice with the above homogenization buffer followed by washes with 3.5 M and 7 M urea added to the homogenization buffer. Centrifugation at 100,000g was performed to separate the supernatant and pellets for each wash. The remaining urea insoluble pellet was taken up in 50 mM TEAB with 5% SDS and protein concentration was measured with BCA assay.

### Tryptic Digest

Membrane pellets of urea insoluble sample (200 µg total protein) were suspended in 5% SDS with 50 mM TEAB and DTT was added to 10 mM before incubation at 56°C for 1 hour to reduce disulfide bonds. Reduced cysteines were alkylated by adding IAA to 20 mM and incubating in the dark at room temperature for 30 minutes. Phosphoric acid was added to 2.5% to insolubilize proteins for efficient trapping. Proteins were taken up in 100 mM TEAB in 90% methanol before transferring 75 µg protein to the S-Trap micro. Trapping solution was centrifuged through the filter at 4,000g for 30 seconds. Trapped proteins were washed three times with 100 mM TEAB in 90% methanol with buffer removed by centrifugation at 4,000g. Proteins were digested by diluting in 50 mM TEAB with 5 µg (1:15) sequencing grade trypsin added and incubated for 1.5 hours at 47°C. To elute digested peptides, four sequential elution solutions of 50 mM TEAB, 0.2% aqueous formic acid, 50% ACN in water and 0.2% aqueous formic acid were added and centrifuged through the filter at 1,000g. Eluted peptides from each sample were pooled and dried with vacuum centrifugation. Peptides were taken up in 0.2% aqueous formic acid and stored at -80°C.

### Liquid Chromatography

Peptides were analyzed using a Dionex Ultimate 3000 UHPLC coupled to an Exploris 480 tandem mass spectrometer (Thermo Scientific, San Jose, CA) with sample order randomized. An in-house pulled capillary column was created from 75 µm inner diameter fused silica capillary packed with 1.9 µm ReproSil-Pur C18 beads (Dr. Maisch, Ammerbuch, Germany) to a length of 250 mm. Solvent A was 0.1% aqueous formic acid and solvent B was 0.1% formic acid in acetonitrile. Peptides were loaded (cortex: 200 ng, outer nucleus: 300 ng, inner nucleus: 300 ng) and separated at a flow rate of 200 nL/min on a 95-minute gradient from 2 to 29% B, followed by a 14-minute washing gradient and 35-minute blank injection between runs. The exact gradient was determined by linearized separation of the top 50% most intense cortical peptide signals by Gradient Optimization Analysis Tool (GOAT 1.0.1).^22^ Full details of chromatographic gradient are provided in Supplemental Table 1.

### Mass Spectrometry

For DDA injections, the Thermo Exploris 480 was set to acquire in top-20 configuration with auto dynamic exclusion. Precursor spectra were collected from 400 to 1600 m/z at 60,000 resolution (AGC target 3e6, max IIT of 50 ms). MS/MS spectra were collected on peptidic precursors between charge state +2 and +5 achieving a minimum AGC of 1e4. MS/MS scans were collected at 15,000 resolution (AGC target of 1e5, max IIT of 35 ms) with an isolation width of 1.6m/z. For DIA injections the Exploris 480 instrument was configured to acquire 35×20m/z (400-1100 m/z) precursor isolation window DIA spectra (30,000 resolution, AGC target 1e6, max IIT 55 ms, 27 NCE) using a staggered window pattern with window placements optimized by Skyline.^23^ Precursor spectra (385-1115 m/z, 60,000 resolution, AGC target 3e6, max IIT 100 ms) were interspersed every 25 ms/ms spectra. In all experiments, default charge state was set to +3, S-Lens RF level set at 40%, NCE set at 27 and data collected in profile mode. See Supplemental Table 2 for complete windowing scheme.

### Data Analysis

For analysis of DDA data, RAW files were searched in MaxQuant^24^ (1.6.7.0) without variable modifications, match between runs enabled, and all other settings set as default. For analysis of DIA data, RAW files were converted to mzML files in MSConvert,^25^ with staggered window deconvolution performed to improve precursor specificity. Processed DIA files were searched in DIA-NN^20^ (1.7.16) with an Intel Core i7-7700 CPU at 3.60 GHz utilizing 8 threads. For all searches, up to one missed trypsin cleavage was allowed on peptides 7-30 residues in length with N-terminal M excision and cysteine carbamidomethylation enabled. All fragments between m/z 200 and 1800 were considered. In each search, the neural network classifier was run in double pass mode with likely interferences removed and retention time dependent normalization initiated between technical replicates from the same region of the lens with high accuracy quantitation enabled. An initial search of all files produced a library which was used to search the data a second time (termed match between runs). Precursors and protein groups were filtered at 1% FDR and contaminants in the MaxQuant contaminant list were removed. All library-free DIA and DDA analyses were searched against a UniProt SwissProt canonical human fasta database (UP000005640, downloaded 3/12/2021, 7,657 entries). For the library free search, deep learning-based spectra and retention time prediction was used for library generation. In a separate library free search, one variable deamidation of glutamine or asparagine was also allowed. For searches against the pan-human SWATH atlas, a consensus speclib file (SAL00035) from Rosenberger et al^19^ was downloaded and used for a library-based search in DIA-NN with identical search settings to library free analysis. Comparisons and analyses were initiated through custom R scripts with peptides having >1% q-value in any DIA-NN q calculation omitted.

Intraregional peptide abundance measurements were independently normalized by DIA-NN abundance normalization based on the 40% of peptides with the lowest CV (<10% CV). To improve normalization, trimmed mean of M-values (TMM) normalization was performed within regions. Interregional peptide abundances were then median normalized. For comparisons of identification efficiency, imputation was not done. When ontology changes were considered in DIA data, the R package DMwR was used for knn-imputation with scaled data pre-processing. Peptides were considered for imputation if detected in 2 of 3 intraregional technical replicates. Protein and peptide abundance was calculated by the diann R package function diann_maxlfq available at https://github.com/vdemichev/diann-rpackage. Proteins were only assembled on peptides considered proteotypic. For DDA comparison, raw intensity peptide abundance values were normalized as above and treated with the diann_maxlfq to retain similarity of protein group quantitation.

Gene association data was downloaded from the Uniprot database (04/06/2021). PSEA-quant^26^ was initiated through a Java command line interface with R scripts for data preparation. Volcano plots were prepared with limma moderated p-values calculated in the eb.fit function.^27^ Quantity of overlapping features were defined by unique razor protein identity or peptides with charge state not considered. For quantitative abundance comparison of DDA and DIA, a linear median normalization factor was determined for proteins identified in both modes and applied to all DDA protein groups. Searches with deamidation enabled were normalized as above, with an additional TMM normalization calculation based on unmodified peptides detected in each search scheme. For all statistical enrichment calculations, PANTHER^28^ was used with the 02/01/2021 GO database applying a Fisher’s exact test with FDR calculated. Significance was set at 0.01 for PANTHER searches. All proteins identified in the modification-free DIA-NN search were included as part of the statistical background.

## Results and Discussion

### Benchmarking against DDA

To evaluate the hypothesis that DIA outperforms DDA in lens protein quantitation, we injected three separate regions of the lens in triplicate to the Exploris 480 in both acquisition modes. DDA analysis was performed in MaxQuant and DIA analysis in DIA-NN. For this comparison, a library free DIA search with no variable modifications was used. In total, 2,161 protein groups and 14,222 peptides were identified in DDA experiments. Previously, as many as 1,251 protein groups had been identified in a cohort of lenses.^14^ The improvement in identifications is facilitated in part by the isolation of insoluble proteins and of the Exploris instrument compared to an older Q Exactive instrument. Prior measurements of the lens have shown significant decreases in protein identification quantity within the outer nucleus and inner nucleus regions of the lens due to organelle-associated protein degradation and targeted degradation of proteins that become misfolded with age. The trend of decreasing protein/peptide identifications in progressively older fibers is replicated in the optimized DDA dataset where there is an approximate 300% increase in proteins and peptides (Figure 2A,B).

**Figure 2.**
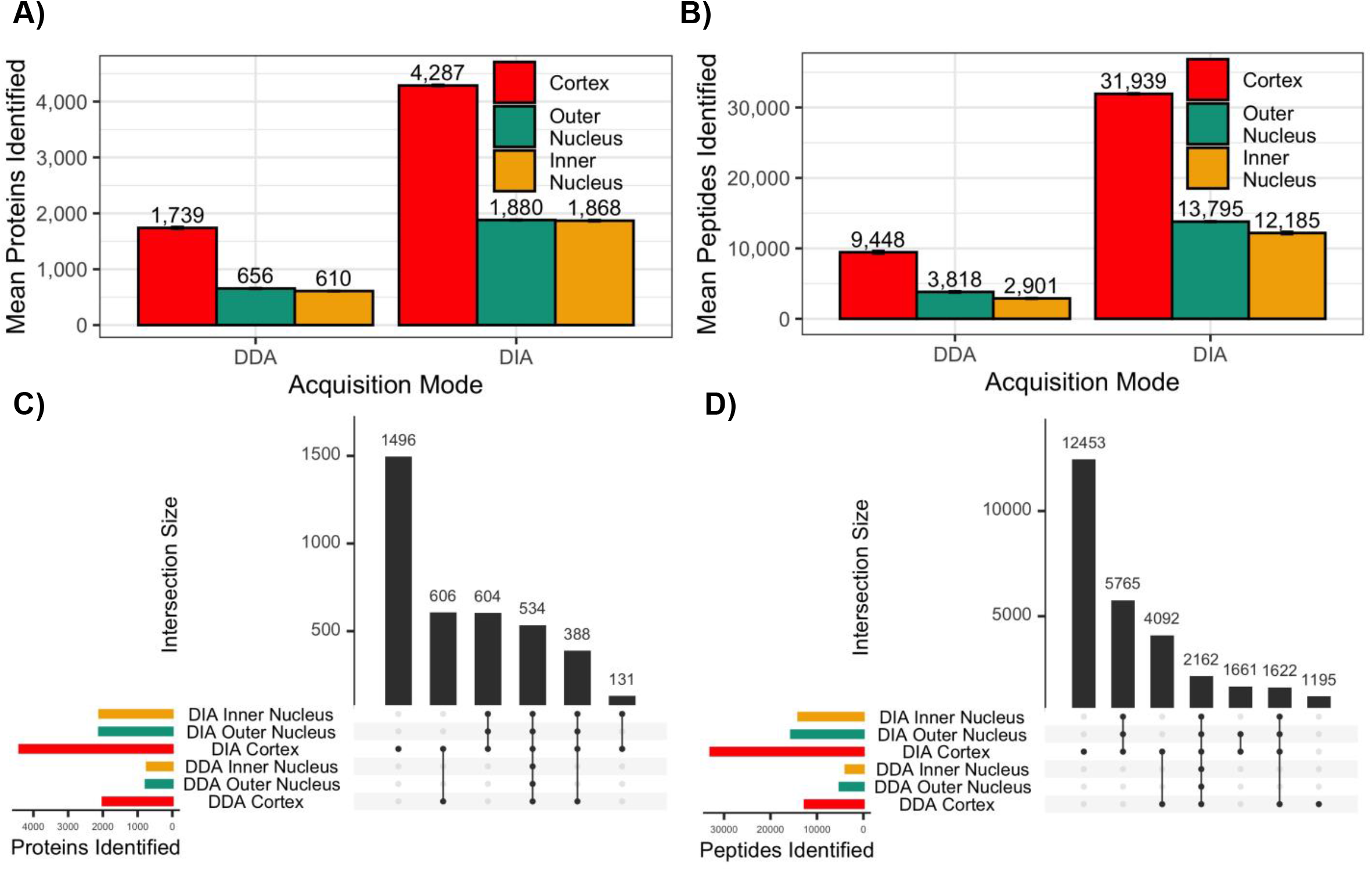
Comparison of Data Dependent (DDA) and Data Independent (DIA) acquisition scan modes for 3 technical replicates of 3 lens regions, PTMs disabled. Only unique leading proteins considered. In total, 4,535 proteins and 35,465 peptides were detected across all experiments. DIA clearly outperforms DDA for identifications and search provides low CV values in protein (A) and peptide (B) comparisons. Upset plots of proteins(C, 82% represented) and peptides (D, 80% represented) reveals near complete coverage of DIA proteome/peptidomes.

Though lens proteomes are less complex than HeLa or other cell culture systems, BRP fractionation established identification of 5,466 proteins in a juvenile mouse lens in 10, 100-minute injections.^29^ In this study, similar proteome coverage was achieved in a single 95-minute gradient where 4,652 protein groups and 33,516 peptides are accessible by single injection DIA experiments, the majority of which are detected in the lens cortex (Figure 2A,B). Juvenile lenses do not undergo significant aging and retain more proteins associated with early lens morphogenesis; therefore, it is not expected that an 18-year-old human lens be more complex than especially young mouse lenses.

Relative to DDA, DIA facilitated a 206% improvement in protein identifications and 235% improvement in total peptide identifications. As with DDA, there is a significant decrease in total identifiable proteins (Figure 2A) as cortical fibers mature to outer nuclear proteins. Unlike DDA results, DIA identification performance of the outer nucleus was nearly matched in the inner nucleus where fibers exceed a decade of age. It is suggested that while still present, a large subset of the nuclear proteome is not detectable with DDA due to biasing limitations. DIA also facilitated greater coverage of identified proteins, with an approximate 7 associated peptides to every protein in DIA, and 5 in DDA. With maturation of the nucleus, the nuclear peptidome, unlike the proteome becomes less populated by unique peptides (Figure 2B). Sequence alignment was not performed on the 1,610 peptides not measured in the inner nucleus; however, the near-complete incorporation of common age-related modifications including deamidation, oxidation, N-and C-terminal truncation may contribute to this difference in searches, like this one, where PTMs are not considered.

To validate the specificity of DIA library free search relative to DDA search algorithms, overlap analysis of leading proteins and peptides was done (Figure 2C,D). Leading proteins were considered to reduce protein group assembly differences between MaxQuant and DIA-NN peptide assignment algorithms. From the 2,161 DDA proteins identified in total, 71 were not included in the DIA dataset, with 38 being identified in an assembled DIA protein group as non-leading or library FDR > 1%. The remaining 33 unmatched proteins (1.5% DDA IDs) show no ontological enrichment against the lens proteome, leading us to determine this discrepancy as insignificant towards protein-network insights. A landmark of success for DIA, 21,243 peptides were uniquely measured with DIA. This figure exceeds the total quantity of peptides identified with DDA, greatly improving the robustness of potential protein quantitation. For proteins measured in DIA and DDA experiments, there is strong regional agreement which further validates the specificity of DIA library free search for peptide assignment. Among peptides, 1,195 from the cortical DDA dataset were not matched to DIA experiments. While not tested, it is expected that the dynamic charge detector facilitated different peptide fragmentation by variable collision energy in DDA whereas DIA employs a static collision energy.

As a consequence of AGC target parameters and chromatographic co-elution of high abundance peptides, peptides of putatively low abundance are not identified in DDA experiments. To demonstrate the distribution of lens protein abundance, we compared the quantitative distribution of calculated protein abundances in each region of the lens (Supplemental Figure 1A,B,C). Search engine quantitation bias was reduced by protein assembly and normalization with the MaxLFQ algorithm in R. For proteins detected in paired studies, a constant necessary to median normalize DDA data to DIA was calculated and applied to all DDA peptides before protein group assembly. For proteins in both DDA and DIA experiments, a non-significant difference of abundance distribution was associated with each regional comparison (Wilcoxon Test). In all regions, the abundance distribution of proteins detected by DDA and DIA had higher means and medians than proteins exclusive to DIA or in the DIA dataset as a whole (p < 2⨯10^−16^). Therefore, we conclude that DIA enables enhanced accessibility to the low abundance lens proteome.

### Evaluating the Use of Pan-Human or Library Free DIA search

Pan-human libraries provide a convenient, sample agnostic approach to DIA assay development. These libraries are highly effective in well-studied tissues or cell lines. Relative to library-free methods, the pan-human library has a significantly decreased search space, decreasing the time needed to process data. To evaluate the utility of pan-human libraries in lens proteomics, we independently searched each sample with and without the pan-human library provided by Rosenberger, Koh, et al.^19^ Since the pan-human library does not have variable modifications annotated, variable modification remained disabled in the library free search. As before, we compared regional feature identifications and performed overlap analysis (Figure 3). Consistent with library free DIA search, the pan-human cortex showed increase protein group diversity relative to the outer and inner nucleus samples. Though much more rapid for data analysis of all 9 replicates (162 minutes for pan-human, 1295 minutes for library free on a personal computer), it is clear that library free searching significantly improves identification quantity of protein groups and peptides. Although the pan-human library search identified more proteins and peptides (3,699/26,147 respective mean IDs in the cortex) than DDA, there is significant variation in nuclear peptide assignment (Figure 3B). In the outer nucleus, CV amounts to 8.6% of mean identifications, and 7.7% in the inner nucleus within technical replicates. Discrepancy is likely caused by a limited quantity of transitions used to define precursors but may be due to false matches.

**Figure 3.**
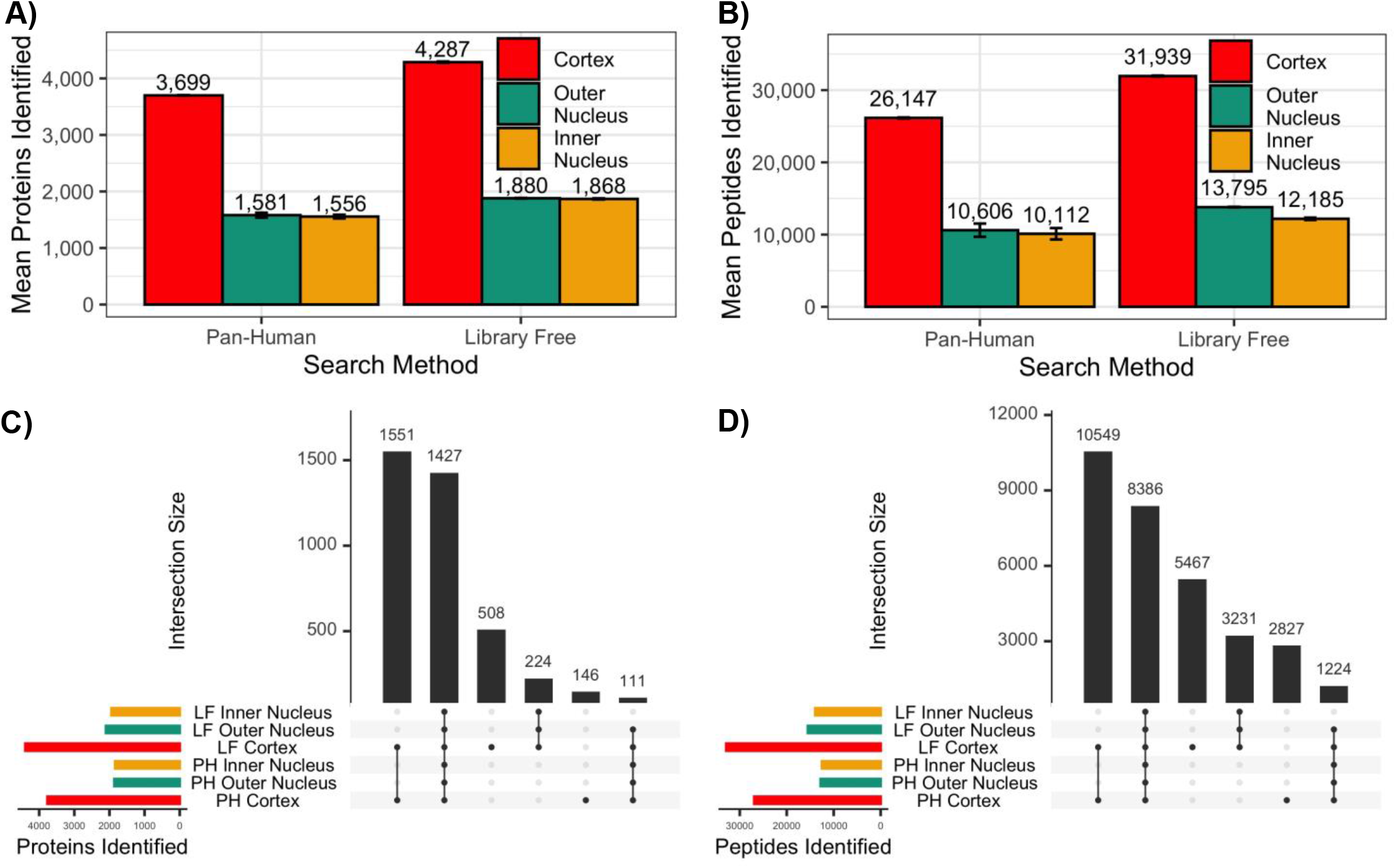
Comparison of Pan-Human (PH) library search with Library Free (LF) DIA-NN Searching, PTMs disabled. Only unique leading proteins considered for identification. A) Protein identifications in each lens region, B) Peptide identifications in each lens region; no modifications were considered. Library-free search produces a greater quantity of IDs filtered at 1% FDR. Upset plots of protein groups (C, 85% represented) and peptides (D, 83% represented) identified in at least one technical replicate of each search method.

In overlap analysis (Figure 3C,D), it was clear that both pan-human and library free methods effectively identify the majority of peptides and proteins expressed in all regions of the lens. However, a significant quantity of proteins (508) and peptides (5,467) were identified in the cortex of the library free search and not in pan-human search. A smaller quantity of cortical features (146 proteins, 2,827 peptides) was uniquely identified in the pan-human search. To determine the biological impact of these unique features, we performed statistical-enrichment analysis of unique library free proteins (859) and pan-human proteins (200) against the lens background (Table 1). No ontologies were statistically enriched from the pan-human search. However, the large quantity of lens-specific ontologies uniquely identified in the library free search demonstrate that the pan-human library does not adequately sample lens fiber cells for annotation of lens biology. Of great significance, Aquaporin-0, one of the most abundant lens proteins, is not in the pan-human library. For future lens studies and studies where PTMs are desirable to measure, it is then suggested that library free search be used in the absence of an exhaustive library.

**Table 1.** Ontological enrichment of proteins detected in library free DIA search and not in Pan-Human DIA search (n = 859). No ontologies were enriched among proteins identified in Pan-Human DIA search and not in library free DIA search (n = 200). Ontology enrichment and statistical analysis performed by PANTHER^28^, using all identified lens proteins as the statistical background.

| GO Term | Annotation | # in Set | Fold Enrichment | FDR |
| --- | --- | --- | --- | --- |
| **Unique in Library Free Search** |  |  |  |  |
| **GOBP** |  |  |  |  |
| GO:0002088 | Lens development in camera-type eye | 26 | 3.56 | 0.0003 |
| GO:0050953 | Sensory perception of light stimulus | 43 | 3.11 | <1x10<sup>-4</sup> |
| GO:0007601 | Visual perception | 43 | 3.11 | <1x10<sup>-4</sup> |
| GO:0043010 | Camera-type eye development | 51 | 2.66 | 0.0006 |
| GO:0007600 | Sensory perception | 52 | 2.30 | <1x10<sup>-4</sup> |
| GO:0001654 | Eye development | 55 | 2.56 | 0.0003 |
| GO:0048880 | Sensory system development | 55 | 2.51 | <1x10<sup>-4</sup> |
| GO:0150063 | Visual system development | 55 | 2.51 | <1x10<sup>-4</sup> |
| GO:0007423 | Sensory organ development | 66 | 2.37 | <1x10<sup>-4</sup> |
| GO:0050877 | Nervous system process | 69 | 1.83 | 0.0024 |
| **GOMF** |  |  |  |  |
| GO:0005212 | Structural constituent of eye lens | 15 | 4.11 | 0.0001 |
| GO:0004888 | Transmembrane signaling receptor activity | 34 | 2.13 | 0.0003 |
| GO:0022857 | Transmembrane transporter activity | 80 | 1.65 | 0.0001 |
| GO:0005215 | Transporter activity | 90 | 1.57 | 0.0002 |
| **GOCC** |  |  |  |  |
| GO:0005877 | Integral component of plasma membrane | 93 | 1.80 | <1x10<sup>-4</sup> |
| GO:0031226 | Intrinsic component of plasma membrane | 101 | 1.78 | <1x10<sup>-4</sup> |
| GO:0016021 | Integral component of membrane | 334 | 1.41 | <1x10<sup>-4</sup> |
| GO:0031224 | Intrinsic component of membrane | 345 | 1.41 | <1x10<sup>-4</sup> |
| GO:0005886 | Plasma membrane | 310 | 1.22 | 0.0060 |
| GO:0071944 | Cell periphery | 351 | 1.21 | 0.0024 |

### Impact of fiber cell aging on proteome composition

A result of aging, deamidation of asparagine or glutamine is the most prevalent PTM in the lens.^30^ While deamidation may be initiated by glutaminase-like enzymes, non-enzymatic deamidation is most common in the lens.^31,32^ To investigate changes in the lens proteome as a function of fiber cell age, we included one variable deamidation in the library free search parameters. Inclusion of a variable deamidation greatly increased the search space over a modification restricted search and may negatively impact FDR calculations. As a result, search time increased from 1,295 minutes with no modification enabled to 3,619 minutes with one variable deamidation allowed: calculations performed on a personal computer. Here, search time increased linearly with search space. In total, 4,564 protein groups and 48,474 peptides were detected. Among these, 1,681 proteins were partially deamidated at 5,577 unique deamidation sites. All of these figures exceed the capabilities of previous human lens proteomic measurements. In this study, we have not considered the peptide-centric effect that deamidation has on lens fiber aging, but rather included these peptides in protein assembly. While possible that some peptides are only identified as deamidated, the functional implication of site-specific modifications are beyond the scope of this discussion.

To evaluate the progressive changes in proteome composition and putative function as a function of fiber cell age, we first evaluated feature overlap between regions (Figure 4A,B). Regional segmentation of the lens allows clear comparisons of fiber cell aging whereas whole-lens measurements previously published are most representative of the high protein density cortical fibers. Exceptionally, 1,844 protein groups (41% of all identified) and 16,250 peptides (34%) were detected as common among lens regions. Here, DIA allows more comparisons in each region than proteins measured in any previous human lens DDA experiment. Thus, changes in whole-protein networks as a function of fiber cell age are best determined with DIA.

**Figure 4.**
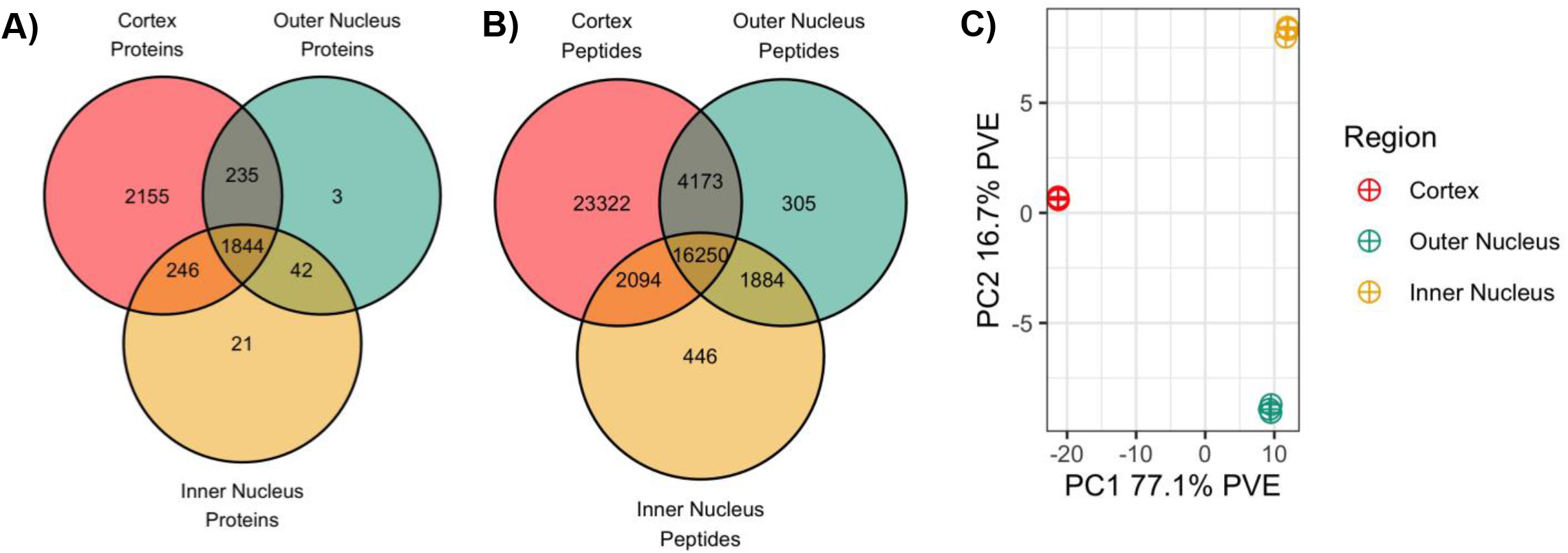
The spatial distribution and abundance of peptides and proteins across the lens displays significant differences as a function of fiber cell age. Data with one variable deamidation used for analysis, intact protein groups used for comparisons. A) Lens protein group overlap with one technical replicate required for consideration, B) Lens peptide overlap, charge states combined, one technical replicate required for consideration. C) PCA demonstrated that cortical fiber and nuclear fiber compositions show significant differences on PC axis 1, and abundance of nuclear fiber proteins contributes to separation on PC axis 2.

It is expected that few or no unique proteins are found in the inner nucleus relative to the outer nucleus, however 265 inner nucleus proteins were detected in the inner nucleus and not in the outer nucleus (Figure 4A). In reference to the lens proteome, enriched parent terms are generally translation related (nuclear-transcribed mRNA catabolic process, nonsense-mediated decay, SRP-dependent co-translational protein targeting to the membrane, viral transcription, translation initiation, rRNA processing, structural constituent of ribosome, cytosolic large ribosomal subunit). It is not expected that these modules, in the absence organelles, are active. It is possible that proteins unique to the inner nucleus are part of lens morphogenesis and are not degraded during early lens development. Nonetheless, in the absence of organelles, it is unlikely that these proteins are metabolically active, and these changes should be considered non-functional. For the 238 proteins detected in the outer nucleus and not in the inner nucleus, no significant protein-network level enrichment was detected. For the 2,155 proteins measured in the cortex but not in either nuclear region, there is no significant functional enrichment in reference to the lens background, but cellular component ontologies including ER membrane, cytosol and integral component of the membrane are enriched in this protein subset.

After numeric comparisons of identification, we investigated protein abundance related differences in each region. With the 1,844 proteins identified in all regions, principal component analysis was performed (Figure 4C). The majority of variance in the dataset is explained by the transition of fiber cells from more classical cellular function in cortical fibers, to the less metabolically active fibers in nuclear regions of the lens, separating cortical and nuclear fibers along PC1. The 50 most descriptive cortical loadings are associated with extracellular matrix assembly, ATP:ADP antiporter activity and basement membrane formation while nuclear loadings are most correlated to cortical cytoskeleton organization and structural constituent of eye lens (data not shown). Secondarily, proteins in the inner nucleus and outer nucleus separated along axis 2. The top 50 PC2 loadings correlated with the outer nucleus are weakly enriched for GDP/GTP binding and GTPase activity. There is weak enrichment of structural constituent of eye lens and phosphatase activity ontologies in loadings most associated with the inner nucleus. While these findings are consistent with previous lens biology studies, they do not take full advantage of the rich DIA dataset. Ontology searches based on a small subset of differentially expressed proteins may be too conservative to identify changes associated with low abundance proteins and analyses considering all identified proteins without considering associated abundance may increase type 1 error rate. Therefore, we set out to perform ontology analysis sensitive to each protein and its associated abundance.

It is clear from pairwise comparisons of protein expression (Figure 5A,B) that the density of most proteins in the cortex exceeds that of either the outer or inner nucleus. As a validation of quantitative comparisons, there is a measured significant increase in γ-crystallin density in each nuclear region respective to the cortex. Unlike the majority of other lens proteins, γ-crystallin is sustained at higher quantities in the lens nucleus due to its role in lens morphogenesis and establishment of the lens gradient of refractive index.^33^ In contrast to previous discussion of insignificant change between the outer nucleus and inner nucleus of the lens, an insignificant, but appreciable trend emerges in the outer nucleus wherein outer nucleus proteins have higher abundance than inner nuclear proteins, which is expected with gradual protein degradation (Figure 5C). In the young lens, the maturation differences between the inner and outer nucleus are proportionally more significant than in an older lens. We suspect that with age, the moderate change in protein abundance between nuclear regions will approach homogeneity except for those proteins differentially expressed by biological function or spatially targeted cellular stress.

**Figure 5.**
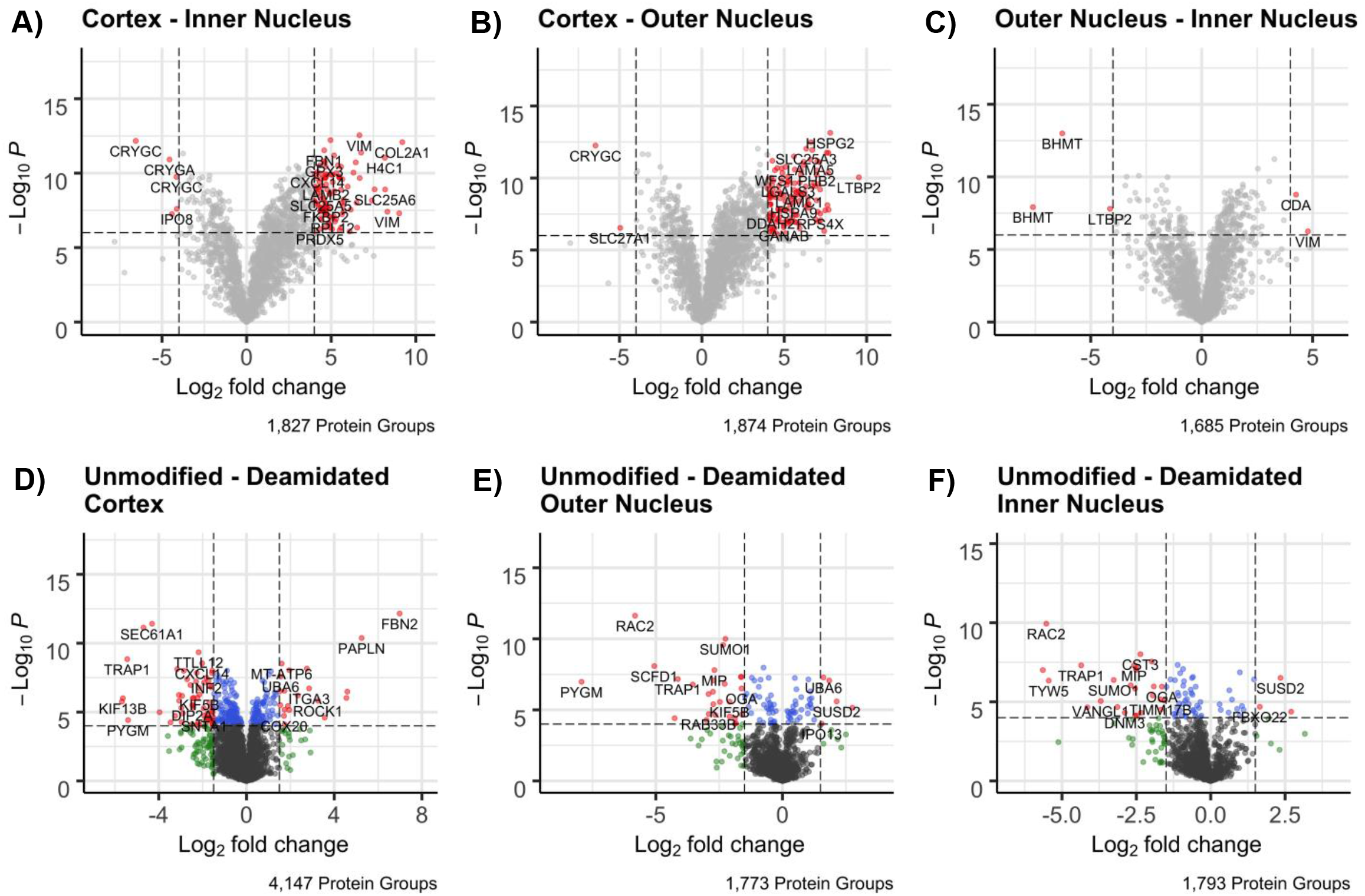
Comparison of protein abundances in the lens. A-C) Regional comparisons of the deamidation enabled search. Limma moderated p-value cutoff was set to 1×10^−6^ with a log_2_FC significance level of 4. Multiple annotation for the same protein may be present because only the leading protein gene association is shown. Full annotation of protein groups is in supplemental table 3. Only values which were detected in all 3 replicates and not imputed were used for comparison. Cortical and outer nuclear proteins tend towards higher abundance over inner nucleus, but the changes between cortical and nuclear fibers is a more discrete change than outer nucleus-inner nucleus. D-F) Protein intensity of compared deamidation disabled and enabled datasets. Statistical null set at 0 Log_2_FC. Proteins with low abundance deamidated peptides are more likely to shift to a positive Log_2_FC. Nonetheless, there is a moderate trend for proteins to have negative FC. In cortical fibers, there is a low degree of expected deamidation. These fibers may be newly formed or be many months old. In the outer nucleus, there is a greater transition of proteins towards the left (deamidation) side of the plot. This trend is best seen in the inner nucleus where proteins may be older than a decade. Few proteins remain at higher intensity in the non-deamidated form. Full annotation of this data is not shown due to less quantitative stringency of comparisons.

To evaluate changes in biology at a protein-network level, abundance-linked ontology analysis was performed. Crystallin proteins make up a disproportionate part of the proteome, so comparisons based on binary expression/non-expression do not represent the dynamic range of the lens proteome. Instead, a rank-based comparison was performed in PSEA-Quant – a modification of the GSEA algorithm.^26^ As may be expected, each region was independently enriched for lens ontologies such as visual perception (GO:0007601) and lens fiber cell development (GO:0070307). Several ontologies were independently enriched in each region but are better delineated in between region calculations.

Between region calculations suffer from the same limitations as previously mentioned; the dramatic dynamic range of the lens proteome abundances does not warrant binary enriched/non-enriched ontology analysis. To quantify changes while between region abundance changes, we again employed PSEA-quant. Briefly, the ratio of protein abundances was computed between each combination of technical replicates of two regions. Only leading proteins that were detected or imputed in all samples were used (n = 1,954). The rank-based enrichment method then identifies changes that are linked to changes between samples, with additional weighting given to proteins with significant changes. For our discussion here, we limit ontological enrichment comparisons to those between the cortex and inner nucleus.

In total, 133 ontologies were enriched in cortex in relation to the inner nucleus, and 73 of the inverse relationship (Table 2, Supplemental Table 4). While it is to be expected that organelle specific proteins are more abundant in the outer cortex relative to the inner nucleus, several inferences related to MCS function can be made. First, there is enrichment of cell-to-cell junctions in young fibers, inferring gradual age-related degradation of extracellular transport networks, even in a young lens, consistent with the formation of a barrier between the cortex and outer nucleus.^4,34^ Next, in the absence of a complex proteome, proteins in the inner nucleus are best suited for oxidoreductase activity by glutathione conjugation. In this model, oxidized glutathione is reduced by glutathione reductase, yielding NADP+ which can be reduced and used by glucose 6-phosphate dehydrogenase (G6PDH) to convert glucose-6-phosphate to 6-phosphogluconate as part of the pentose phosphate pathway. Previous results in an aged rat lens demonstrated that G6PDH is catalytically inactive in the inner nucleus^35^ leaving fibers devoid of glycolysis function, even though the protein machinery is present.^36^

**Table 2.** Reduced table of ontologies specifically enriched in the cortex or inner nucleus over respective partner. Less specific, redundant and terms less relevant to lens biology were filtered. Full list of significant annotations found in Supplemental Table 4. Calculations performed with PSEA-quant.^26^

| GO Term | Annotation | PES | p-value | Proteins in Set |
| --- | --- | --- | --- | --- |
| **Enriched in Cortex over Inner Nucleus** |  |  |  |  |
| GO:0000184 | Nuclear-transcribed mRNA catabolic process, nonsense-mediated decay | 42.48 | <1.0x10<sup>-4</sup> | 54 |
| GO:0022625 | Cytosolic large ribosomal subunit | 22.62 | <1.0x10<sup>-4</sup> | 27 |
| GO:0031982 | Vesicle | 115.25 | <1.0x10<sup>-4</sup> | 171 |
| GO:0005886 | Plasma membrane | 233.86 | <1.0x10<sup>-4</sup> | 363 |
| GO:0006414 | Translational elongation | 40.01 | <1.0x10<sup>-4</sup> | 49 |
| GO:0005912 | Adherens junction | 34.49 | <1.0x10<sup>-4</sup> | 47 |
| GO:0043624 | Cellular protein complex disassembly | 40.06 | <1.0x10<sup>-4</sup> | 50 |
| GO:0006886 | Intracellular protein transport | 118.86 | <1.0x10<sup>-4</sup> | 176 |
| GO:0070161 | Anchoring junction | 38.60 | <1.0x10<sup>-4</sup> | 53 |
| GO:0006415 | Translational termination | 38.29 | <1.0x10<sup>-4</sup> | 46 |
| GO:0015075 | Ion transmembrane transporter activity | 39.36 | <1.0x10<sup>-4</sup> | 53 |
| GO:0016071 | mRNA metabolic process | 95.93 | <1.0x10<sup>-4</sup> | 144 |
| GO:0031090 | Organelle membrane | 222.46 | <1.0x10<sup>-4</sup> | 347 |
| GO:0032984 | Protein-containing complex disassembly | 41.32 | <1.0x10<sup>-4</sup> | 52 |
| GO:0005544 | Calcium-dependent phospholipid binding | 9.07 | 0.0002 | 11 |
| GO:0008324 | Cation transmembrane transporter activity | 26.73 | 0.0002 | 37 |
| GO:0045296 | Cadherin binding | 8.91 | 0.0008 | 10 |
| GO:0015077 | Monovalent inorganic cation transmembrane transporter activity | 19.90 | 0.0008 | 27 |
| KEGG | ECM receptor interaction | 15.31 | <1.0x10<sup>-4</sup> | 17 |
| KEGG | Ribosome | 38.05 | <1.0x10<sup>-4</sup> | 46 |
| KEGG | Focal adhesion | 35.02 | 0.0001 | 49 |
| Reactome | Translation | 75.19 | 0.0003 | 112 |
| Reactome | Metabolism of amino acids and derivatives | 86.99 | 0.0004 | 132 |
| Reactome | L1CAM interactions | 28.91 | 0.001 | 41 |
| **Enriched in Inner Nucleus over Cortex** |  |  |  |  |
| GO:0007601 | Visual perception | 19.74 | <1.0x10<sup>-4</sup> | 33 |
| GO:0005212 | Structural constituent of eye lens | 12.84 | <1.0x10<sup>-4</sup> | 17 |
| GO:0016765 | Transferase activity, transferring alkyl or aryl (other than methyl) groups | 9.38 | <1.0x10<sup>-4</sup> | 15 |
| GO:0019318 | Hexose metabolic process | 32.67 | <1.0x10<sup>-4</sup> | 62 |
| GO:0007600 | Sensory perception | 28.03 | <1.0x10<sup>-4</sup> | 51 |
| GO:0002088 | Lens development in camera-type eye | 8.34 | <1.0x10<sup>-4</sup> | 12 |
| GO:0016491 | Oxidoreductase activity | 60.55 | 0.0001 | 120 |
| GO:0046983 | Protein dimerization activity | 71.11 | 0.0001 | 141 |
| GO:0097354 | Prenylation | 3.53 | 0.0001 | 5 |
| GO:0008318 | Protein prenyltransferase activity | 3.53 | 0.0001 | 5 |
| GO:0045762 | Positive regulation of adenylate cyclase activity | 3.51 | 0.0002 | 5 |
| GO:0016056 | Rhodopsin mediated signaling pathway | 4.54 | 0.0002 | 7 |
| GO:0016614 | Oxidoreductase activity, acting on CH-OH group of donors | 17.62 | 0.0003 | 33 |
| GO:0006749 | Glutathione metabolic process | 9.19 | 0.0003 | 16 |
| GO:0016616 | Oxidoreductase activity, acting on the CH-OH group of donors, NAD or NADP as acceptor | 17.62 | 0.0005 | 33 |
| GO:0008238 | Exopeptidase activity | 14.65 | 0.0006 | 27 |
| GO:0006096 | Glycolytic process | 11.22 | 0.0006 | 20 |
| GO:0006733 | Oxidoreduction coenzyme metabolic process | 12.75 | 0.0008 | 24 |
| GO:0046872 | Metal ion binding | 172.67 | 0.0009 | 355 |
| GO:0006656 | Phosphatidylcholine biosynthetic process | 2.17 | 0.001 | 3 |
| KEGG | Glutathione metabolism | 11.72 | 0.0006 | 21 |
| KEGG | Glycolysis gluconeogenesis | 14.76 | 0.0008 | 27 |
| Reactome | Metabolism of carbohydrates | 37.91 | <1.0x10<sup>-4</sup> | 73 |
| Reactome | Beta catenin independent WNT signaling | 27.14 | 0.0005 | 53 |
| Reactome | PCP CE pathway | 23.13 | 0.0007 | 45 |
| Reactome | Fructose metabolism | 2.65 | 0.0008 | 4 |
| Reactome | Biological oxidations | 16.75 | 0.0009 | 32 |

A benefit of greater proteome coverage is that a greater proportion of networks may be considered for ontological significance. In PSEA-quant, networks that meet p-value and FDR significance fail to be deemed significant when <3 proteins from the studied ontology are present. For ontology networks enriched in the inner nucleus (Supplemental Table 4, Table 2), 22 of the significant ontologies are represented by less than 10 proteins in the dataset. Indeed, when DDA data was processed through the PSEA-quant pipeline with identical treatment, only 61 ontologies were enriched in either the cortex or inner nucleus (data not shown) – all with redundancy to the 206 ontologies from DIA results. Among cortically enriched DIA ontologies, few are sparsely populated as a result of the significant improvement in identifications. Thus, the enhanced proteome coverage improves preliminary identification of protein networks that are influential in lens fiber aging. Lastly, the enhanced peptidome coverage afforded by DIA provides a more robust quantitative assembly of protein groups, leading to more reproducible and biologically representative protein abundance measurement.

Finally, we evaluated the impact of fiber cell age-related deamidation on total age-related protein abundance in the lens. Peptide level search results with deamidation enabled and disabled were TMM normalized before protein assembly and significance testing (Figure 5D,E,F). In both the outer cortex and outer nucleus, the vast majority of the proteome is not highly deamidated, while the inner nucleus shows a trend towards deamidation. In the relatively young 18-year-old lens, it is not expected that the majority of the lens would be deamidated, but the gradual accumulation of deamidation demonstrated here establishes specificity of library-free search towards deamidated peptides. The young lens is less age- effected than an older lens but sustains deamidation on 1,681 proteins. The majority of these proteins are believed to be deamidated non-enzymatically, and prior DDA evidence (Schey, unpublished data) suggests that there will be even more peptides deamidated as a function of age. Given the assumption of non-enzymatic deamidation, ontological assignment of differences is not relevant until a greater quantity of biological replicates are available.

## Conclusion

Here, we have shown that the use of DIA methodology for lens proteomics greatly improves identification of low abundance proteins and have demonstrated an effective way to search the data with lens biology in mind. The gradual shift towards protein deamidation and proteome specialization for oxidoreductase activity clearly demonstrates that this method is sensitive to age-related fiber maturation measurements – and the enhanced ability of DIA to measure low abundance peptides improves protein-network level characterization of lens aging. Additionally, the application of PSEA-quant here demonstrates that an ontological search that is sensitive to protein abundance is well suited towards assessing lens biology. It is reasonable to suggest that the improved protein network coverage afforded by DIA paired to sensitive analysis pipelines may allow improved insights into human lens aging and cataract formation.

## Supporting information

Supplemental File 4 DIA Deamidation Report

Supplemental File 3 DIA Pan Human Report

Supplemental File 2 DIA No Modifications Report

Supplemental File 8 DIA Deamidation Protein Groups

Supplemental File 7 DIA Pan Human Protein Groups

Supplemental File 6 DIA No Modifications Protein Groups

Supplemental File 5 DDA Protein Groups

Supplemental File 1 DDA Evidence

## Data Availability

Reports from DIA-NN, MaxQuant evidence file and normalized protein group assemblies are available as Supplemental Files 1-8.

## Acknowledgements

We would like to acknowledge the Vanderbilt Mass Spectrometry Research Center, particularly Kristie L. Rose and W. Hayes McDonald for assistance in developing DIA methods for this experiment. We also thank Zhen Wang and all other members of the Schey lab for thoughtful conversations in the development of this work. Finally, we acknowledge Vadim Demichev for his ongoing support of DIA data analysis. Financial support is acknowledged from NIH grants EY013462 (KLS), EY008126, and T32 GM065086 (LSC).

## Supplemental Figures

**Supplemental Figure 1.**
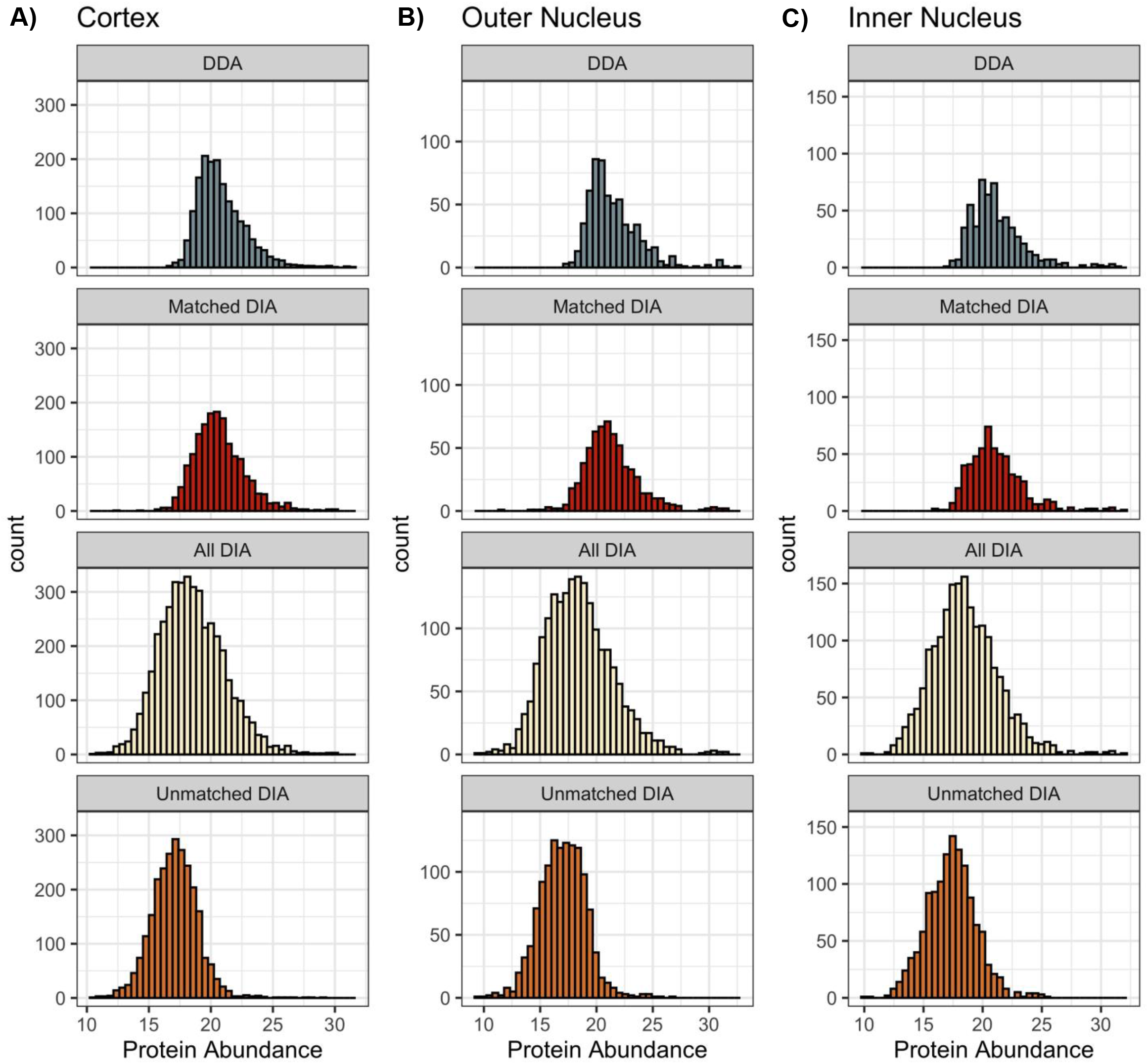
Histogram comparison of Log2 abundances between DDA and DIA, each calculated with the diann R package. For each region of the lens A) Cortex, B) Outer Nucleus, C) Inner Nucleus, quantitative median normalization factors were calculated for matched protein groups between DDA and DIA and applied to all unmatched protein groups. DDA abundances (grey) visually resemble the distribution of matched ID DIA proteins (red). The majority of proteins uniquely identified in DIA (orange) were of lower mean abundance compared to the DDA proteome. The resulting complete DIA experiment (beige) has a moderately decreased median and wider quantitative distribution than DDA.

**Supplemental Table 1.** Full chromatography details with gradient calculated at 3-minute intervals by GOAT^22^ with an assumed 11-minute retention time delay.

| **Time (min)** | **%B** |
| --- | --- |
| 0.0 | 2.0 |
| 1.0 | 2.0 |
| 2.0 | 2.0 |
| 3.0 | 2.0 |
| 6.0 | 7.1 |
| 9.0 | 8.4 |
| 12.0 | 9.4 |
| 15.0 | 10.1 |
| 18.0 | 11.1 |
| 21.0 | 11.8 |
| 24.0 | 12.3 |
| 27.0 | 12.6 |
| 30.0 | 13.7 |
| 33.0 | 14.4 |
| 36.0 | 14.7 |
| 39.0 | 15.0 |
| 42.0 | 15.5 |
| 45.0 | 16.2 |
| 48.0 | 16.5 |
| 51.0 | 17.0 |
| 54.0 | 17.7 |
| 57.0 | 18.4 |
| 60.0 | 18.7 |
| 63.0 | 19.2 |
| 66.0 | 19.9 |
| 69.0 | 20.6 |
| 72.0 | 21.5 |
| 75.0 | 22.4 |
| 78.0 | 24.1 |
| 81.0 | 90.0 |
| 83.0 | 90.0 |
| 84.0 | 2.0 |
| 85.0-95.0 | 2.0 |

**Supplemental Table 2.** Calculated mass windows for DIA experiments. Optimization done in Skyline^23^ from 400-1100 m/z with staggered window deconvolution selected. Windows exceeding the specified range were discarded.

| Calculated m/z Window |  |
| --- | --- |
| 400.4319-420.441 | 410.4364-430.4455 |
| 420.441-440.4501 | 430.4455-450.4546 |
| 440.4501-460.4592 | 450.4546-470.4637 |
| 460.4592-480.4683 | 470.4637-490.4728 |
| 480.4683-500.4774 | 490.4728-510.4819 |
| 500.4774-520.4865 | 510.4819-530.491 |
| 520.4865-540.4956 | 530.491-550.5001 |
| 540.4956-560.5047 | 550.5001-570.5092 |
| 560.5047-580.5138 | 570.5092-590.5183 |
| 580.5138-600.5229 | 590.5183-610.5274 |
| 600.5229-620.5319 | 610.5274-630.5365 |
| 620.5319-640.541 | 630.5365-650.5456 |
| 640.541-660.5501 | 650.5456-670.5547 |
| 660.5501-680.5592 | 670.5547-690.5638 |
| 680.5592-700.5683 | 690.5638-710.5729 |
| 700.5683-720.5774 | 710.5729-730.582 |
| 720.5774-740.5865 | 730.582-750.5911 |
| 740.5865-760.5956 | 750.5911-770.6002 |
| 760.5956-780.6047 | 770.6002-790.6093 |
| 780.6047-800.6138 | 790.6093-810.6183 |
| 800.6138-820.6229 | 810.6183-830.6274 |
| 820.6229-840.632 | 830.6274-850.6365 |
| 840.632-860.6411 | 850.6365-870.6456 |
| 860.6411-880.6502 | 870.6456-890.6547 |
| 880.6502-900.6593 | 890.6547-910.6638 |
| 900.6593-920.6684 | 910.6638-930.6729 |
| 920.6684-940.6775 | 930.6729-950.682 |
| 940.6775-960.6866 | 950.682-970.6911 |
| 960.6866-980.6957 | 970.6911-990.7002 |
| 980.6957-1000.7048 | 990.7002-1010.7093 |
| 1000.7048-1020.7138 | 1010.7093-1030.7184 |
| 1020.7138-1040.7229 | 1030.7184-1050.7275 |
| 1040.7229-1060.732 | 1050.7275-1070.7366 |
| 1060.732-1080.7411 | 1070.7366-1090.7457 |
| 1080.7411-1100.7502 |  |

**Supplemental Table 3.**
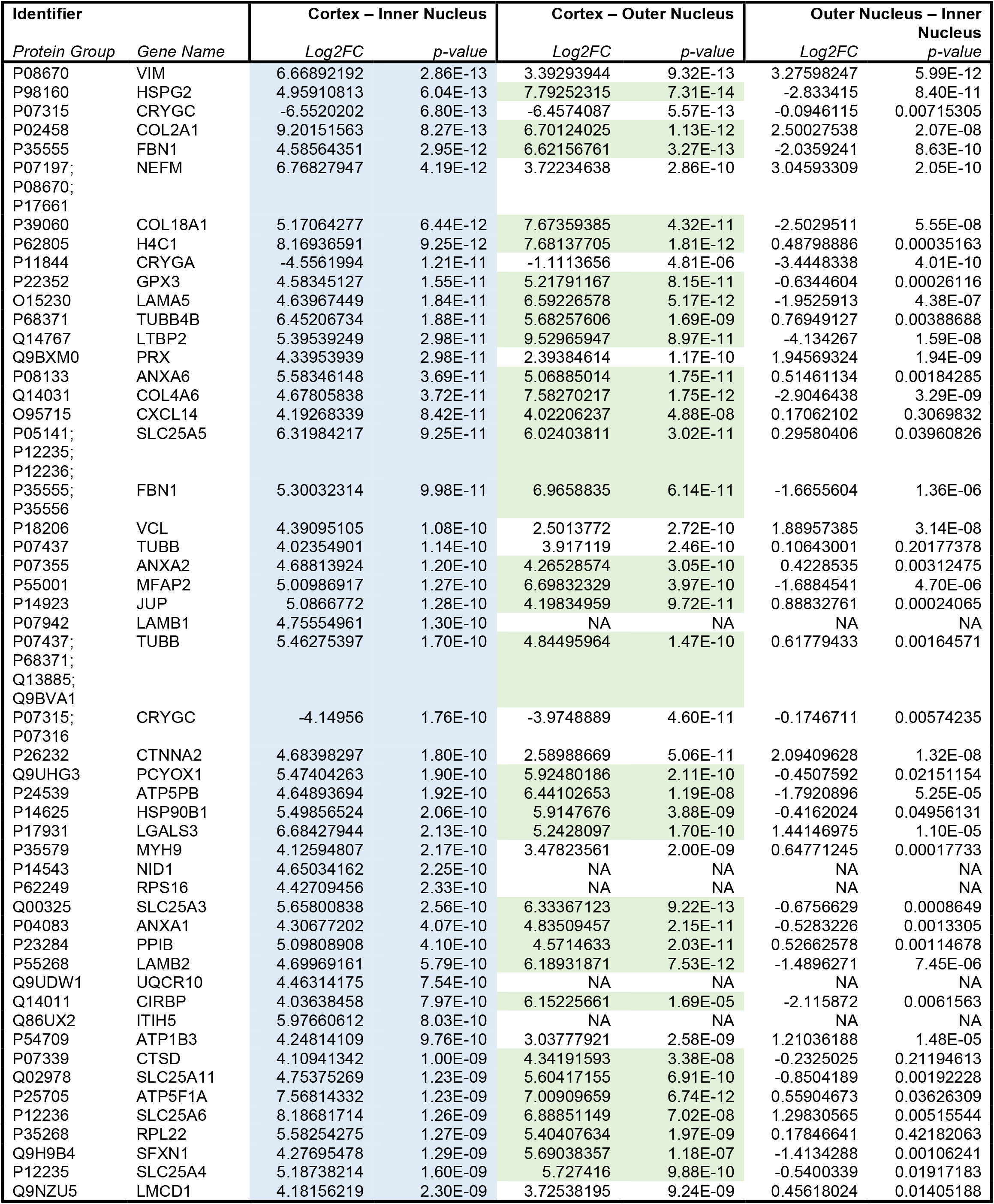
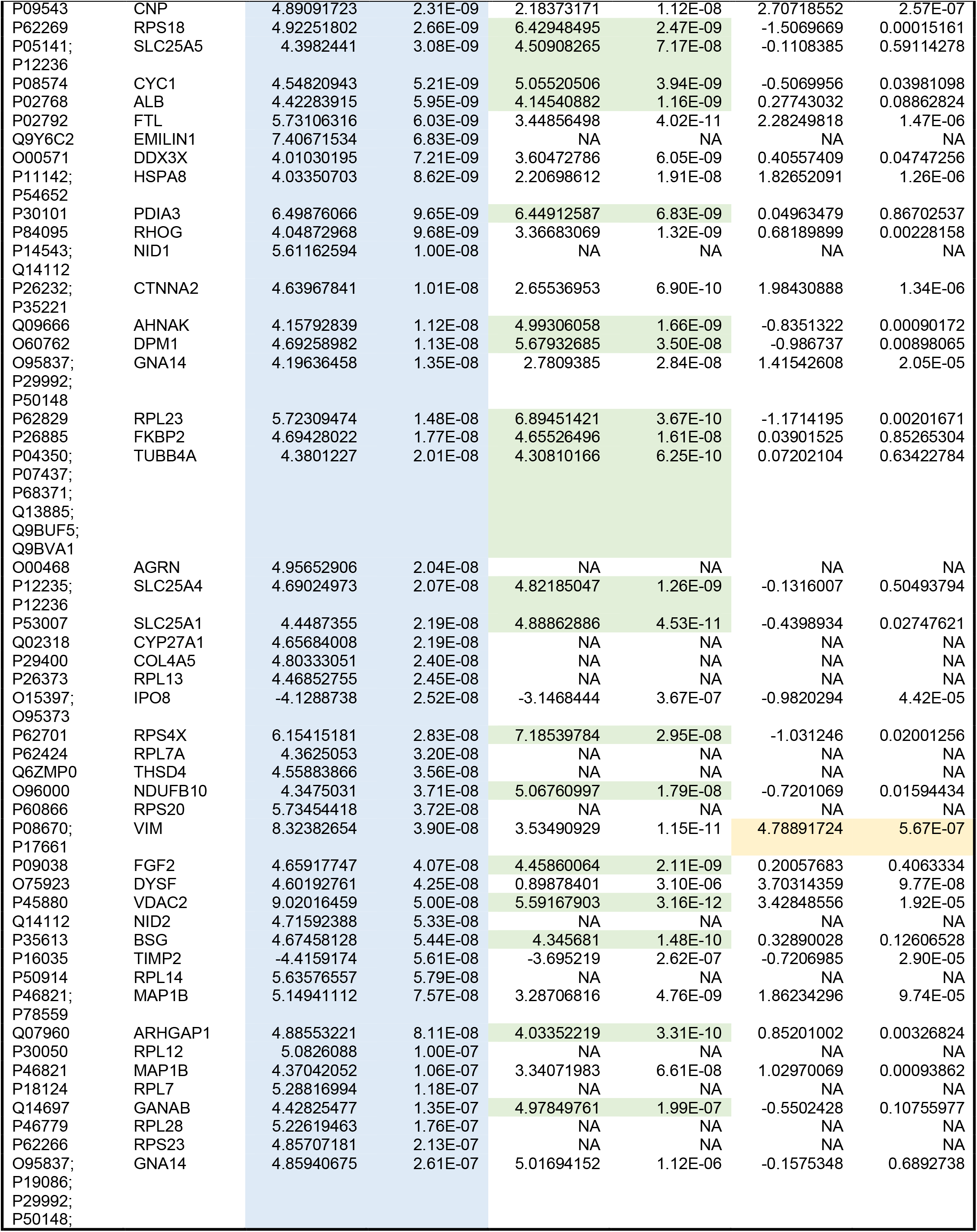
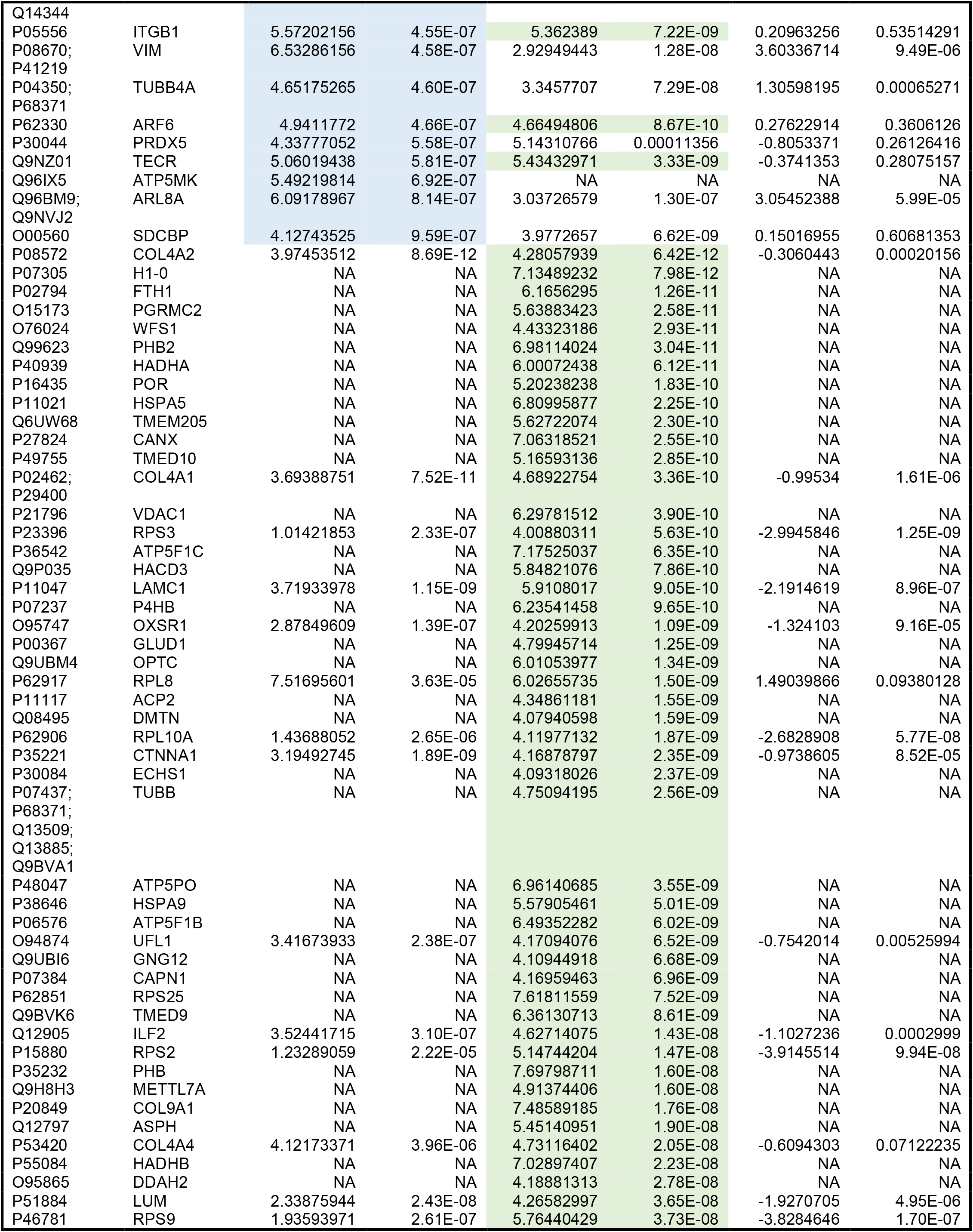
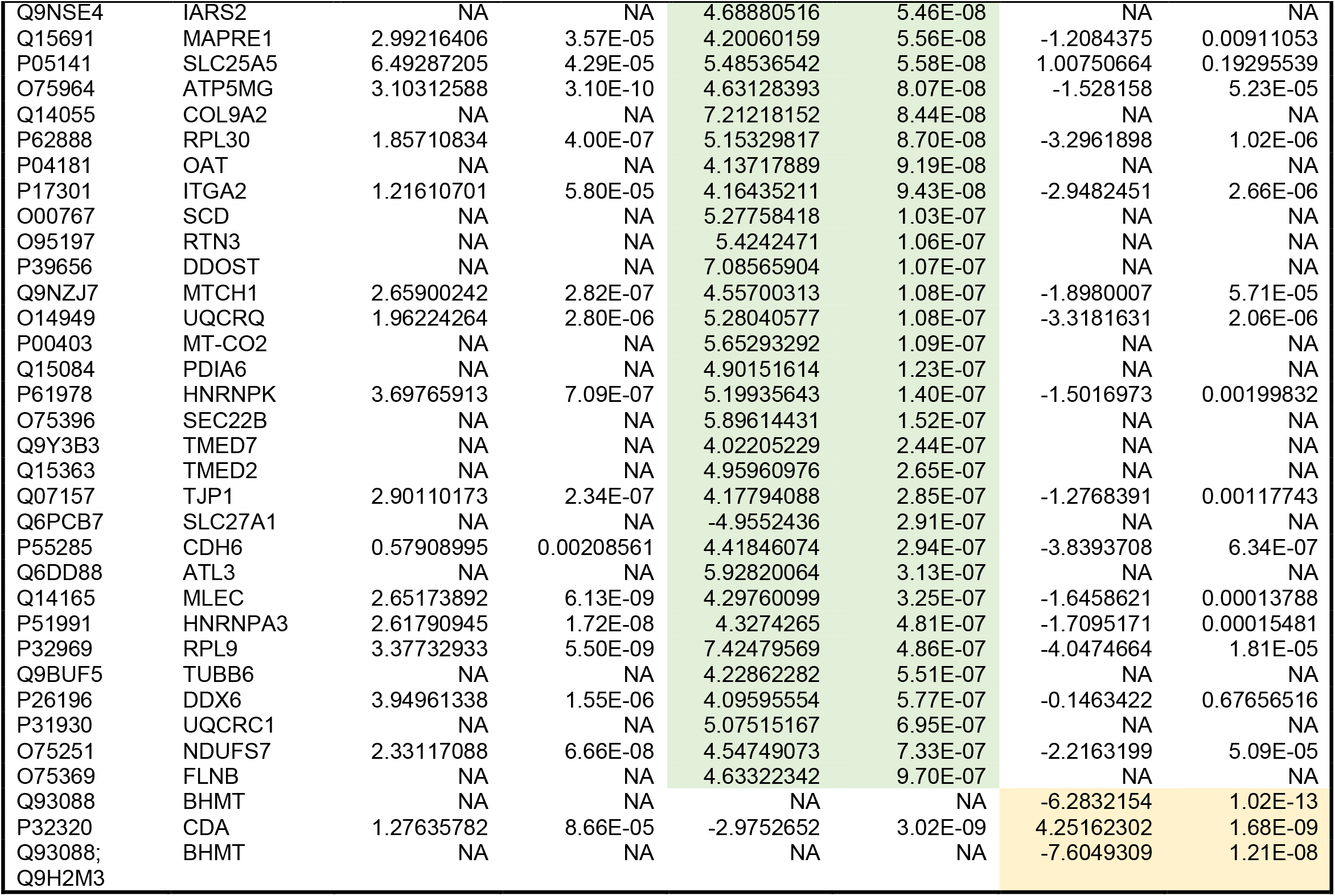
Complete protein group and gene group ID of each protein identified as differentially enriched in supplemental figure 2 (n=190). Only proteins with limma moderated p-values below 1E-6 and Log2 FC in excess of 4 are shown.

**Supplemental Table 4.** Complete list of ontologies (n=206) enriched in either the cortex or inner nucleus over the other region. Ontology calculations performed by PSEA-quant^26^ with CV tolerance 0.5, 10,000 iterations used for p-value calculation.

| GO Term | Annotation | PES | p-value | Proteins in Set |
| --- | --- | --- | --- | --- |
| **Enriched in Cortex over Inner Nucleus** |  |  |  |  |
| GO:0031974 | Membrane-enclosed lumen | 70.5365 | <1.0x10<sup>-4</sup> | 95 |
| GO:0005604 | Basement membrane | 15.54 | <1.0x10<sup>-4</sup> | 19 |
| GO:0006612 | Protein targeting to membrane | 51.185 | <1.0x10<sup>-4</sup> | 62 |
| GO:0000184 | Nuclear-transcribed mrna catabolic process, nonsense-mediated decay | 42.485 | <1.0x10<sup>-4</sup> | 54 |
| GO:0022625 | Cytosolic large ribosomal subunit | 22.6225 | <1.0x10<sup>-4</sup> | 27 |
| GO:0034655 | Nucleobase-containing compound catabolic process | 110.8015 | <1.0x10<sup>-4</sup> | 161 |
| GO:0006605 | Protein targeting | 65.0145 | <1.0x10<sup>-4</sup> | 86 |
| GO:0071702 | Organic substance transport | 225.7785 | <1.0x10<sup>-4</sup> | 351 |
| GO:0031982 | Vesicle | 115.25 | <1.0x10<sup>-4</sup> | 171 |
| GO:0051234 | Establishment of localization | 326.6375 | <1.0x10<sup>-4</sup> | 509 |
| GO:0005886 | Plasma membrane | 233.866 | <1.0x10<sup>-4</sup> | 363 |
| GO:0006414 | Translational elongation | 40.015 | <1.0x10<sup>-4</sup> | 49 |
| GO:0043062 | Extracellular structure organization | 43.002 | <1.0x10<sup>-4</sup> | 55 |
| GO:0005759 | Mitochondrial matrix | 27.1465 | <1.0x10<sup>-4</sup> | 36 |
| GO:0000956 | Nuclear-transcribed mRNA catabolic process | 47.017 | <1.0x10<sup>-4</sup> | 63 |
| GO:0043933 | Protein-containing complex subunit organization | 160.473 | <1.0x10<sup>-4</sup> | 246 |
| GO:0005912 | Adherens junction | 34.4925 | <1.0x10<sup>-4</sup> | 47 |
| GO:0043624 | Cellular protein complex disassembly | 40.069 | <1.0x10<sup>-4</sup> | 50 |
| GO:0015935 | Small ribosomal subunit | 17.7745 | <1.0x10<sup>-4</sup> | 22 |
| GO:0016043 | Cellular component organization | 360.0055 | <1.0x10<sup>-4</sup> | 569 |
| GO:0019866 | Organelle inner membrane | 46.838 | <1.0x10<sup>-4</sup> | 63 |
| GO:0044391 | Ribosomal subunit | 41.923 | <1.0x10<sup>-4</sup> | 52 |
| GO:0031012 | Extracellular matrix | 33.3745 | <1.0x10<sup>-4</sup> | 45 |
| GO:0046700 | Heterocycle catabolic process | 115.055 | <1.0x10<sup>-4</sup> | 169 |
| GO:0005789 | Endoplasmic reticulum membrane | 39.407 | <1.0x10<sup>-4</sup> | 52 |
| GO:0019083 | Viral transcription | 37.7415 | <1.0x10<sup>-4</sup> | 45 |
| GO:0005200 | Structural constituent of cytoskeleton | 22.6465 | <1.0x10<sup>-4</sup> | 30 |
| GO:0046907 | Intracellular transport | 177.6925 | <1.0x10<sup>-4</sup> | 269 |
| GO:0044270 | Cellular nitrogen compound catabolic process | 114.819 | <1.0x10<sup>-4</sup> | 169 |
| GO:0006886 | Intracellular protein transport | 118.8685 | <1.0x10<sup>-4</sup> | 176 |
| GO:0070161 | Anchoring junction | 38.602 | <1.0x10<sup>-4</sup> | 53 |
| GO:0071840 | Cellular component organization or biogenesis | 361.837 | <1.0x10<sup>-4</sup> | 573 |
| GO:0006415 | Translational termination | 38.2925 | <1.0x10<sup>-4</sup> | 46 |
| GO:0072594 | Establishment of protein localization to organelle | 63.31 | <1.0x10<sup>-4</sup> | 87 |
| GO:1901361 | Organic cyclic compound catabolic process | 120.529 | <1.0x10<sup>-4</sup> | 179 |
| GO:0030574 | Collagen catabolic process | 10.9265 | <1.0x10<sup>-4</sup> | 13 |
| GO:0005198 | Structural molecule activity | 116.5215 | <1.0x10<sup>-4</sup> | 160 |
| GO:0006613 | Cotranslational protein targeting to membrane | 41.004 | <1.0x10<sup>-4</sup> | 49 |
| GO:0045184 | Establishment of protein localization | 188.9685 | <1.0x10<sup>-4</sup> | 291 |
| GO:0042470 | Melanosome | 38.7975 | <1.0x10<sup>-4</sup> | 50 |
| GO:0006810 | Transport | 322.4605 | <1.0x10<sup>-4</sup> | 503 |
| GO:0022411 | Cellular component disassembly | 68.8565 | <1.0x10<sup>-4</sup> | 92 |
| GO:0070062 | Extracellular exosome | 32.9365 | <1.0x10<sup>-4</sup> | 44 |
| GO:0022627 | Cytosolic small ribosomal subunit | 16.119 | <1.0x10<sup>-4</sup> | 19 |
| GO:0006402 | mRNA catabolic process | 48.1525 | <1.0x10<sup>-4</sup> | 65 |
| GO:0015075 | Ion transmembrane transporter activity | 39.3665 | <1.0x10<sup>-4</sup> | 53 |
| GO:0015031 | Protein transport | 185.297 | <1.0x10<sup>-4</sup> | 286 |
| GO:0003735 | Structural constituent of ribosome | 41.793 | <1.0x10<sup>-4</sup> | 54 |
| GO:0072599 | Establishment of protein localization to endoplasmic reticulum | 41.5735 | <1.0x10<sup>-4</sup> | 50 |
| GO:0019439 | Aromatic compound catabolic process | 116.7855 | <1.0x10<sup>-4</sup> | 173 |
| GO:0032963 | Collagen metabolic process | 10.9265 | <1.0x10<sup>-4</sup> | 13 |
| GO:0016020 | Membrane | 421.909 | <1.0x10<sup>-4</sup> | 674 |
| GO:0070013 | Intracellular organelle lumen | 56.1445 | <1.0x10<sup>-4</sup> | 72 |
| GO:0005201 | Extracellular matrix structural constituent | 13.0005 | <1.0x10<sup>-4</sup> | 14 |
| GO:0065010 | Extracellular membrane-bounded organelle | 32.9365 | <1.0x10<sup>-4</sup> | 44 |
| GO:0016482 | Cytosolic transport | 92.9645 | <1.0x10<sup>-4</sup> | 135 |
| GO:0016071 | mRNA metabolic process | 95.9325 | <1.0x10<sup>-4</sup> | 144 |
| GO:0031090 | Organelle membrane | 222.463 | <1.0x10<sup>-4</sup> | 347 |
| GO:0032984 | Protein-containing complex disassembly | 41.322 | <1.0x10<sup>-4</sup> | 52 |
| GO:0022617 | Extracellular matrix disassembly | 11.734 | <1.0x10<sup>-4</sup> | 14 |
| GO:0045047 | Protein targeting to ER | 41.5735 | <1.0x10<sup>-4</sup> | 50 |
| GO:0006413 | Translational initiation | 52.02 | <1.0x10<sup>-4</sup> | 72 |
| GO:0048770 | Pigment granule | 38.7975 | <1.0x10<sup>-4</sup> | 50 |
| GO:0005788 | Endoplasmic reticulum lumen | 17.189 | <1.0x10<sup>-4</sup> | 19 |
| GO:0043230 | Extracellular organelle | 32.9365 | <1.0x10<sup>-4</sup> | 44 |
| GO:0042383 | Sarcolemma | 11.763 | 0.0001 | 15 |
| GO:0005581 | Collagen trimer | 9.606 | 0.0001 | 10 |
| GO:0006928 | Movement of cell or subcellular component | 122.56 | 0.0001 | 187 |
| GO:0006412 | Translation | 66.6785 | 0.0001 | 98 |
| GO:0031966 | Mitochondrial membrane | 60.38 | 0.0001 | 88 |
| GO:0032774 | RNA biosynthetic process | 89.8865 | 0.0001 | 135 |
| GO:0022857 | Transmembrane transporter activity | 49.8445 | 0.0001 | 71 |
| GO:0005544 | Calcium-dependent phospholipid binding | 9.0785 | 0.0002 | 11 |
| GO:0050839 | Cell adhesion molecule binding | 14.1955 | 0.0002 | 18 |
| GO:0019898 | Extrinsic component of membrane | 20.6575 | 0.0002 | 28 |
| GO:0005911 | Cell-cell junction | 39.452 | 0.0002 | 56 |
| GO:0034329 | Cell junction assembly | 31.771 | 0.0002 | 45 |
| GO:0008324 | Cation transmembrane transporter activity | 26.736 | 0.0002 | 37 |
| GO:0044291 | Cell-cell contact zone | 12.7445 | 0.0003 | 16 |
| GO:0051271 | Negative regulation of cellular component movement | 13.782 | 0.0003 | 18 |
| GO:0031410 | Cytoplasmic vesicle | 86.503 | 0.0003 | 131 |
| GO:0044764 | Multi-organism cellular process | 119.078 | 0.0003 | 183 |
| GO:0014704 | Intercalated disc | 11.04 | 0.0004 | 14 |
| GO:0051235 | Maintenance of location | 15.787 | 0.0004 | 21 |
| GO:0016323 | Basolateral plasma membrane | 21.3535 | 0.0006 | 29 |
| GO:0030054 | Cell junction | 78.201 | 0.0007 | 118 |
| GO:0016032 | Viral process | 119.078 | 0.0007 | 183 |
| GO:0044419 | Interspecies interaction between organisms | 126.0995 | 0.0007 | 194 |
| GO:0044403 | Symbiotic process | 119.078 | 0.0007 | 183 |
| GO:0017111 | Nucleoside-triphosphatase activity | 106.7195 | 0.0007 | 164 |
| GO:0045296 | Cadherin binding | 8.917 | 0.0008 | 10 |
| GO:0015077 | Monovalent inorganic cation transmembrane transporter activity | 19.902 | 0.0008 | 27 |
| GO:0051651 | Maintenance of location in cell | 12.4125 | 0.0009 | 16 |
| KEGG | ECM receptor interaction | 15.3155 | <1.0x10<sup>-4</sup> | 17 |
| KEGG | Ribosome | 38.0505 | <1.0x10<sup>-4</sup> | 46 |
| KEGG | Focal adhesion | 35.026 | 0.0001 | 49 |
| KEGG | Parkinsons disease | 22.586 | 0.0001 | 30 |
| KEGG | Huntingtons disease | 28.949 | 0.0002 | 40 |
| KEGG | Small cell lung cancer | 13.32 | 0.0004 | 17 |
| Reactome | Laminin interactions | 15.4535 | <1.0x10<sup>-4</sup> | 16 |
| Reactome | Nonsense mediated decay nmd | 42.6985 | <1.0x10<sup>-4</sup> | 54 |
| Reactome | Extracellular matrix organization | 43.111 | <1.0x10<sup>-4</sup> | 55 |
| Reactome | Nervous system development | 127.377 | <1.0x10<sup>-4</sup> | 190 |
| Reactome | Influenza infection | 48.068 | <1.0x10<sup>-4</sup> | 63 |
| Reactome | SRP dependent cotranslational protein targeting to membrane | 41.7885 | <1.0x10<sup>-4</sup> | 50 |
| Reactome | Collagen degradation | 9.8835 | <1.0x10<sup>-4</sup> | 11 |
| Reactome | Degradation of the extracellular matrix | 20.3545 | <1.0x10<sup>-4</sup> | 25 |
| Reactome | Response of EIF2AK4 GCN2 to amino acid deficiency | 40.775 | <1.0x10<sup>-4</sup> | 50 |
| Reactome | Regulation of expression of slits and ROBOs | 62.3505 | <1.0x10<sup>-4</sup> | 91 |
| Reactome | ECM proteoglycans | 18.5665 | <1.0x10<sup>-4</sup> | 21 |
| Reactome | Non integrin membrane ECM interactions | 19.3275 | <1.0x10<sup>-4</sup> | 22 |
| Reactome | Eukaryotic translation elongation | 40.2355 | <1.0x10<sup>-4</sup> | 49 |
| Reactome | Integrin cell surface interactions | 18.6175 | <1.0x10<sup>-4</sup> | 21 |
| Reactome | Developmental biology | 141.9425 | <1.0x10<sup>-4</sup> | 212 |
| Reactome | Met activates PTK2 signaling | 7.052 | <1.0x10<sup>-4</sup> | 8 |
| Reactome | Collagen biosynthesis and modifying enzymes | 10.1835 | <1.0x10<sup>-4</sup> | 11 |
| Reactome | Collagen formation | 11.9575 | <1.0x10<sup>-4</sup> | 14 |
| Reactome | Assembly of collagen fibrils and other multimeric structures | 9.6625 | <1.0x10<sup>-4</sup> | 11 |
| Reactome | Eukaryotic translation initiation | 51.365 | <1.0x10<sup>-4</sup> | 70 |
| Reactome | rRNA processing | 43.077 | <1.0x10<sup>-4</sup> | 56 |
| Reactome | Selenoamino acid metabolism | 45.43 | <1.0x10<sup>-4</sup> | 58 |
| Reactome | Binding and uptake of ligands by scavenger receptors | 10.6175 | 0.0001 | 13 |
| Reactome | Molecules associated with elastic fibres | 7.0035 | 0.0001 | 8 |
| Reactome | NCAM1 interactions | 7.92 | 0.0001 | 9 |
| Reactome | Disease | 188.331 | 0.0001 | 291 |
| Reactome | Infectious disease | 130.6035 | 0.0001 | 197 |
| Reactome | Elastic fibre formation | 7.0035 | 0.0002 | 8 |
| Reactome | NCAM signaling for neurite out growth | 14.678 | 0.0002 | 19 |
| Reactome | Smooth muscle contraction | 8.1835 | 0.0003 | 10 |
| Reactome | Translation | 75.192 | 0.0003 | 112 |
| Reactome | Metabolism of amino acids and derivatives | 86.999 | 0.0004 | 132 |
| Reactome | Signaling by met | 14.9195 | 0.0007 | 20 |
| Reactome | L1CAM interactions | 28.918 | 0.001 | 41 |
| **Enriched in Inner Nucleus over Cortex** |  |  |  |  |
| GO:0007601 | Visual perception | 19.743 | <1.0x10<sup>-4</sup> | 33 |
| GO:0003008 | System process | 56.885 | <1.0x10<sup>-4</sup> | 112 |
| GO:0003824 | Catalytic activity | 412.0705 | <1.0x10<sup>-4</sup> | 851 |
| GO:0050877 | Nervous system process | 37.025 | <1.0x10<sup>-4</sup> | 70 |
| GO:0042803 | Protein homodimerization activity | 57.578 | <1.0x10<sup>-4</sup> | 112 |
| GO:0005212 | Structural constituent of eye lens | 12.841 | <1.0x10<sup>-4</sup> | 17 |
| GO:0016765 | Transferase activity, transferring alkyl or aryl (other than methyl) groups | 9.383 | <1.0x10<sup>-4</sup> | 15 |
| GO:0050953 | Sensory perception of light stimulus | 19.743 | <1.0x10<sup>-4</sup> | 33 |
| GO:0019318 | Hexose metabolic process | 32.67 | <1.0x10<sup>-4</sup> | 62 |
| GO:0007600 | Sensory perception | 28.0335 | <1.0x10<sup>-4</sup> | 51 |
| GO:0044281 | Small molecule metabolic process | 238.9625 | <1.0x10<sup>-4</sup> | 491 |
| GO:0002088 | Lens development in camera-type eye | 8.3425 | <1.0x10<sup>-4</sup> | 12 |
| GO:0005996 | Monosaccharide metabolic process | 35.7905 | <1.0x10<sup>-4</sup> | 68 |
| GO:0006006 | Glucose metabolic process | 28.6985 | <1.0x10<sup>-4</sup> | 54 |
| GO:0016052 | Carbohydrate catabolic process | 21.9255 | <1.0x10<sup>-4</sup> | 40 |
| GO:0046365 | Monosaccharide catabolic process | 17.72 | 0.0001 | 32 |
| GO:0019320 | Hexose catabolic process | 16.126 | 0.0001 | 29 |
| GO:1900544 | Positive regulation of purine nucleotide metabolic process | 5.173 | 0.0001 | 8 |
| GO:0006575 | Cellular modified amino acid metabolic process | 21.9645 | 0.0001 | 41 |
| GO:0042398 | Cellular modified amino acid biosynthetic process | 7.661 | 0.0001 | 13 |
| GO:0016491 | Oxidoreductase activity | 60.552 | 0.0001 | 120 |
| GO:0046983 | Protein dimerization activity | 71.1105 | 0.0001 | 141 |
| GO:0043436 | Oxoacid metabolic process | 87.658 | 0.0001 | 176 |
| GO:0019362 | Pyridine nucleotide metabolic process | 11.969 | 0.0001 | 22 |
| GO:0097354 | Prenylation | 3.5375 | 0.0001 | 5 |
| GO:0008318 | Protein prenyltransferase activity | 3.5375 | 0.0001 | 5 |
| GO:0031281 | Positive regulation of cyclase activity | 3.5115 | 0.0002 | 5 |
| GO:0045762 | Positive regulation of adenylate cyclase activity | 3.5115 | 0.0002 | 5 |
| GO:0051349 | Positive regulation of lyase activity | 3.5115 | 0.0002 | 5 |
| GO:0030810 | Positive regulation of nucleotide biosynthetic process | 4.496 | 0.0002 | 7 |
| GO:0045981 | Positive regulation of nucleotide metabolic process | 5.173 | 0.0002 | 8 |
| GO:1900373 | Positive regulation of purine nucleotide biosynthetic process | 4.496 | 0.0002 | 7 |
| GO:0022400 | Regulation of rhodopsin mediated signaling pathway | 4.5435 | 0.0002 | 7 |
| GO:0019752 | Carboxylic acid metabolic process | 82.262 | 0.0002 | 164 |
| GO:0016056 | Rhodopsin mediated signaling pathway | 4.5435 | 0.0002 | 7 |
| GO:0046496 | Nicotinamide nucleotide metabolic process | 11.969 | 0.0002 | 22 |
| GO:0004659 | Prenyltransferase activity | 4.638 | 0.0002 | 7 |
| GO:0016614 | Oxidoreductase activity, acting on CH-OH group of donors | 17.6295 | 0.0003 | 33 |
| GO:0006749 | Glutathione metabolic process | 9.1955 | 0.0003 | 16 |
| GO:0042802 | Identical protein binding | 89.5835 | 0.0003 | 179 |
| GO:0006520 | Cellular amino acid metabolic process | 52.6775 | 0.0003 | 103 |
| GO:0006007 | Glucose catabolic process | 13.9 | 0.0004 | 25 |
| GO:0008652 | Cellular amino acid biosynthetic process | 17.2205 | 0.0004 | 32 |
| GO:0016616 | Oxidoreductase activity, acting on the CH-OH group of donors, NAD or NADP as acceptor | 17.6295 | 0.0005 | 33 |
| GO:0006082 | Organic acid metabolic process | 89.6605 | 0.0005 | 180 |
| GO:0018342 | Protein prenylation | 3.5375 | 0.0005 | 5 |
| GO:0008238 | Exopeptidase activity | 14.657 | 0.0006 | 27 |
| GO:0006790 | Sulfur compound metabolic process | 26.116 | 0.0006 | 50 |
| GO:0006096 | Glycolytic process | 11.2235 | 0.0006 | 20 |
| GO:0005615 | Extracellular space | 43.2595 | 0.0006 | 85 |
| GO:0044238 | Primary metabolic process | 467.2315 | 0.0006 | 976 |
| GO:0043603 | Cellular amide metabolic process | 19.016 | 0.0008 | 36 |
| GO:0006733 | Oxidoreduction coenzyme metabolic process | 12.754 | 0.0008 | 24 |
| GO:0072524 | Pyridine-containing compound metabolic process | 13.153 | 0.0008 | 25 |
| GO:0071482 | Cellular response to light stimulus | 6.559 | 0.0008 | 11 |
| GO:0031279 | Regulation of cyclase activity | 4.904 | 0.0009 | 8 |
| GO:0090330 | Regulation of platelet aggregation | 2.164 | 0.0009 | 3 |
| GO:0006081 | Cellular aldehyde metabolic process | 8.882 | 0.0009 | 16 |
| GO:0046872 | Metal ion binding | 172.6725 | 0.0009 | 355 |
| GO:0006656 | Phosphatidylcholine biosynthetic process | 2.1765 | 0.001 | 3 |
| KEGG | Glutathione metabolism | 11.721 | 0.0006 | 21 |
| KEGG | Glycolysis gluconeogenesis | 14.7665 | 0.0008 | 27 |
| Reactome | Metabolism of carbohydrates | 37.9195 | $<1.0 \times 10^{-4}$ | 73 |
| Reactome | TP53 regulates transcription of cell death genes | 5.4785 | 0.0001 | 9 |
| Reactome | The phototransduction cascade | 4.5435 | 0.0001 | 7 |
| Reactome | Glycolysis | 14.465 | 0.0003 | 27 |
| Reactome | ROBO receptors bind AKAP5 | 2.771 | 0.0003 | 4 |
| Reactome | Signaling by WNT | 35.3235 | 0.0004 | 70 |
| Reactome | Beta catenin independent WNT signaling | 27.1485 | 0.0005 | 53 |
| Reactome | PCP CE pathway | 23.138 | 0.0007 | 45 |
| Reactome | Fructose metabolism | 2.654 | 0.0008 | 4 |
| Reactome | Biological oxidations | 16.75 | 0.0009 | 32 |
| Reactome | Transcriptional regulation of TP53 | 26.5975 | 0.0009 | 52 |

## Notes

### Competing Interest Statement

The authors have declared no competing interest.

